# Ultra-sensitive nanoLC-MS of sub nanogram protein samples using second generation micro pillar array LC technology with Orbitrap Exploris 480 and FAIMS PRO

**DOI:** 10.1101/2021.02.10.430648

**Authors:** Karel Stejskal, Jeff Op de Beeck, Gerhard Dürnberger, Paul Jacobs, Karl Mechtler

**Author notes:** KS and JB will contribute equal to the manuscript. **Corresponding Author** Karl Mechtler - IMBA - Institute of Molecular Biotechnology of the Austrian Academy of Sciences, Dr. Bohr Gasse 3, A-1030 Vienna, Austria, IMP - Institute of Molecular Pathology, Campus-Vienna-Biocenter 1, A-1030 Vienna, Austria, Gregor Mendel Institute of Molecular Plant Biology of the Austrian Academy of Sciences, Dr. Bohr Gasse 3, A-1030 Vienna, Austria.

## Abstract

In the light of the ongoing single-cell revolution, scientific disciplines are combining forces to retrieve as much relevant data as possible from trace amounts of biological material. For single cell proteomics, this implies optimizing the entire workflow from initial cell isolation down to sample preparation, liquid chromatography (LC) separation, mass spectrometer (MS) data acquisition and data analysis. To demonstrate the potential for single cell and limited sample proteomics, we report on a series of benchmarking experiments where we combine LC separation on a new generation of micro pillar array columns with state-of-the-art Orbitrap MS/MS detection and High-Field Asymmetric Waveform Ion Mobility Spectrometry (FAIMS). This dedicated limited sample column has a reduced cross section and micro pillar dimensions that have been further downscaled (inter pillar distance and pillar diameter by a factor of 2), resulting in improved chromatography at reduced void times. A dilution series of a HeLa tryptic digest (5-0.05 ng/μL) was used to explore the sensitivity that can be achieved. Comparative processing of the MS/MS data with Sequest HT, MS Amanda, Mascot and SpectroMine pointed out the benefits of using Sequest HT together with INFERYS when analyzing sample amounts below 1 ng. 2855 protein groups were identified from just 1 ng of HeLa tryptic digest hereby increasing detection sensitivity as compared to a previous contribution by a factor well above 10. By successfully identifying 1486 protein groups from as little as 250 pg of HeLa tryptic digest, we demonstrate outstanding sensitivity with great promise for use in limited sample proteomics workflows.

## INTRODUCTION

During the last few years, the sensitivity of LC-MS/MS instrumentation has evolved to a level where consistent identification and quantification of proteins from single cells has become feasible ^1–5^. As opposed to LC-MS/MS analysis of proteins originating from bulk cell populations, proteome analysis of a single or a few selected cells allows attributing biological characteristics to individual cells. It can also provide crucial and unbiased insights in the role that specific cell types play in biological processes ^6^. As already observed in bulk proteomics samples, recent studies demonstrate modest correlation between transcriptomic and proteomic expression levels obtained from single cells ^1,7^. This highlights the complementary nature and pertinence of LC-MS/MS based proteome profiling at the single cell level.

In contrast to single cell analysis at the genomic and transcriptomic level, single cell proteome analysis cannot rely on techniques that allow amplifying trace amounts of protein ^8^. Therefore the entire proteomics workflow needs to be optimized carefully to achieve ultrasensitive analysis and near loss-less processing of protein samples ^9–11^. Recent breakthroughs in the field of low input LC-MS/MS based proteomics have been obtained in parallel by several independent groups, each using a unique approach to tackle the challenges associated with comprehensive single cell proteome analysis ^1,3,12–15^. The key aspects to the successful implementation seem to be a combination of miniaturized and automated sample preparation methods with ultrasensitive LC-MS/MS analysis performed at very low LC flow rates (≤ 100 nL/min), additional ion mobility separation and the latest generation of hybrid or tribrid MS-MS instruments. As from the inception of nanoelectrospray in the 90s by Mann et al. ^16^, the major leaps in sensitivity that can be achieved by combining true nanoliter per minute flow rates with electrospray ionization have been a driving force in the practice of MS based proteomics. However, robust operation at very low flow rates (≤ 100 nL/min) is still a big challenge and requires either highly specialized LC systems that are not widely available, or custom column configurations.

Building further upon a previous contribution where we reported on the benefits of using perfectly ordered micro pillar array based nano HPLC columns for low-input proteomics ^17^, we now present an ultrasensitive LC-MS/MS based proteomics workflow at standard nano flow rates (250 nL/min). By combining outstanding chromatographic performance of a novel dedicated micro pillar array column type with High-Field Asymmetric Waveform Ion Mobility Spectrometry (FAIMS) Pro technology, the latest generation of Orbitrap MS systems and optimized search strategies, we manage to achieve more than a 10 fold increase in detection sensitivity on the protein level as compared to Stadlmann et al. in 2019 ^17^. The potential of this approach for single cell proteome profiling is demonstrated by analyzing a dilution series of HeLa cell tryptic digest from 5 ng down to 50 pg, resulting in consistent identification of 2855 protein groups from 1 ng of HeLa tryptic digest.

## EXPERIMENTAL

To investigate the potential gain in sensitivity that can be achieved for low input proteome profiling, the micro pillar array column was operated with an Ultimate™ 3000 RSLCnano system with ProFlow™ technology (Thermo Fisher Scientific) and coupled to an Orbitrap Exploris™ 480 mass spectrometer (Thermo Fisher Scientific) equipped with a FAIMS pro interface (Thermo Fisher Scientific). The dedicated limited sample micro pillar array column (PharmaFluidics) has a total length of 50 cm and is filled with 2.5 μm diameter non porous silicon pillars that have been positioned at a distance of 1.25 μm in an equilateral triangular grid (Supplementary information figure 1). The stationary phase morphology has been optimized to deliver maximal performance for low input reversed phase liquid chromatography, which is discussed more extensively in the supporting Information.

**Figure 1.**
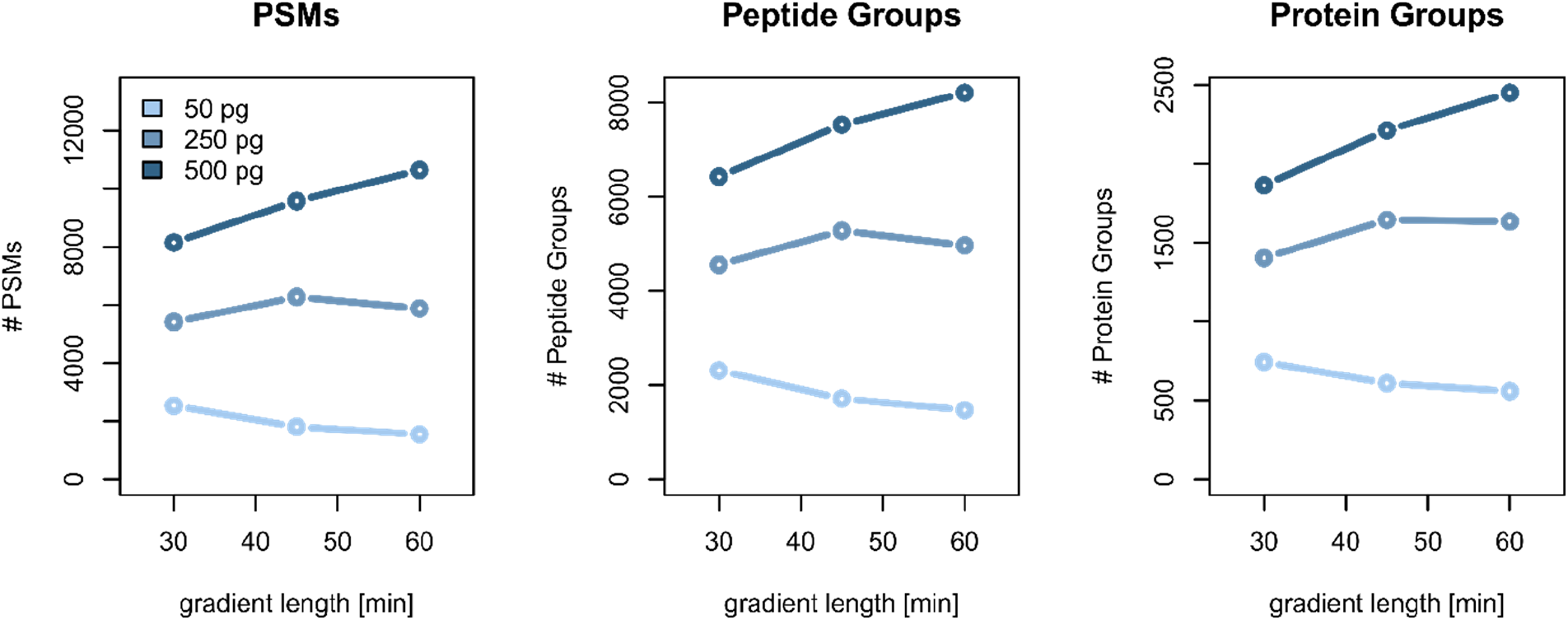
Effect of solvent gradient length on identifications (PSM, Peptide and Protein group level). 50, 250 and 500 pg of HeLa tryptic digest was separated using a non-linear solvent gradient of 30, 45 and 60 minutes.

### Sample preparation

Thermo Scientific™ Pierce™ HeLa Protein Digest Standard (P/N 88328) was obtained from Thermo Scientific™ in lyophilized form and used as test sample for all measurements. 20 μg of lyophilized peptide material in glass vial was reconstituted to a concentration of 100 ng/μL in LC/MS grade water with 0.1% (v/v) formic acid (FA). For the final dilution to the required concentration, peptide solution was diluted directly in autosampler vials with a glass insert (Thermo Scientific™, P/N 500212). The solvent used to perform dilutions was 0.001% (w/v) polyethylene glycol (PEG 20000, Sigma Aldrich, P/N 83100) in LC/MS grade water with 0.1% (v/v) formic acid (FA). To investigate the effect of PEG addition to the sample stability, samples were diluted to 250 pg/μl in autosampler vial with glass insert. Injection was performed over a period of 24 h after dilution at 4 h intervals. A HeLa dilution series was finally prepared with PEG in the sample solvent and samples were injected as technical quadruplicates, starting with the lowest concentration (0.25 ng/μl) and ending with the highest concentration (5 ng/μL).

### LC configuration

The column was placed in the column oven compartment and maintained at a constant temperature of 40°C throughout the entire experiment. A shortened fused silica capillary (20 μm ID x 360 μm OD, length 40 cm, P/N 6041.5293, Thermo Fisher Scientific) was used to connect the column inlet to the Ultimate™ 3000 RSLCnano autosampler injection valve, configured to perform direct injection of 1μL volume sample plugs (1μL sample loop – full loop injection mode). Separation was achieved with stepped linear solvent gradients, all performed at a fixed flow rate of 250 nL/min with various durations of 30, 45 and 60 minutes. Organic modifier content (Acetonitrile acidified with 0.1% v/v formic acid) was first increased from 0.8 to 16% in 24.5, 35.75 and 47 minutes, then increased from 16 to 28% in respectively 7.5, 11.25 and 15 minutes and finally ramped from 28 to 78% in 5 minutes. Mobile phase composition was kept at high organic (78% Acetonitrile acidified with 0.1% v/v formic acid) for 5 minutes to wash the column after which column re-equilibration was performed at low organic (0.8% Acetonitrile acidified with 0.1% v/v formic acid) for 17 minutes.

Efficient transfer of peptides from the LC column to the mass spectrometer was achieved by connecting the column outlet union to a PepSep sprayer 1 (PepSep, P/N PSS1) equipped with 10 μm ID fused silica electrospray emitter with integrated liquid junction (PepSep, P/N PSFSELJ10) using a custom made fused silica capillary with nanoConnect (PepSep, 20 μm ID x 360 μm OD, length 15 cm). A grounded connection was provided between the column outlet union and the grounding pin at the back of the RSLCnano system to prevent electrical current leaking to the LC column. An electrospray voltage of 2.3 kV was applied at the integrated liquid junction of the emitter which was installed on a Nanospray Flex™ ion source (Thermo Fisher Scientific).

### MS Acquisition

The mass spectrometer was operated in positive mode with the FAIMS Pro interface. Compensation voltage was set at −50 V to remove singly charged ions. Data dependent acquisition (DDA) was performed using the following parameters. MS1 resolution was set at 120k with a normalized AGC target 300 % (3e6), a maximum inject time was set to Auto and a scan range of 375 to 1200 m/z. For MS2, resolution was set at 60k with a normalized AGC target of 75 % (7.5e4), maximum inject time of 118 ms. Top 10 abundant precursors (charge state 2-5) within an isolation window of 2 m/z were considered for MS/MS analysis. Dynamic exclusion was set at 120s, mass tolerance of ±10 ppm was allowed, and the precursor intensity threshold was set at 5e3. For precursor fragmentation in HCD mode, a normalized collision energy of 30% was used. Whereas technical replicates were not evaluated for initial gradient length and sample stability optimization, technical quadruplicates were run for the in-depth analysis of the HeLa tryptic digest dilution series (5 to 0.25 ng).

### Data analysis

MS/MS spectra from raw data acquired at various concentrations of HeLa tryptic digest were imported to Proteome Discoverer (PD) (version 2.5.0.400, Thermo Scientific). All database search engines were operated with identical parameter settings. First MS/MS spectra were recalibrated in the PD node “Spectrum Files RC” using the human SwissProt database (*Homo sapiens*; release 2019_06, 20339 sequences, 11360750 residues) and a database of common contaminants (372 sequences, 145187 residues). Recalibration was performed for fully tryptic peptides applying an initial precursor tolerance of 20 ppm and a fragment tolerance of 0.1 Da. Carbamidomethylation of cysteine was set as a fixed modification in the recalibration step. Database searches were performed using the same FASTA databases as described above. Trypsin was specified as proteolytic enzyme, cleaving after lysine (K) and arginine (R) except when followed by proline (P) and up to one missed cleavage was considered. Mass tolerance was limited to 10 ppm at the precursor and 0.02 Da at the fragment level. Carbamidomethylation of cysteine (C) was set as fixed modification and oxidation of methionine (M) as well as loss of methionine at the protein N-terminus were set as variable modifications. Identified spectra were rescored using Percolator ^18^ as implemented in PD and filtered for 1% FDR at the peptide spectrum match and peptide level. Based on this set of common search parameters several database search engines were evaluated for their performance of identifying spectra from low sample amounts, namely MS Amanda ^19^, Sequest HT ^20^ and Mascot 2.2 ^21^. MS Amanda and Sequest HT were evaluated with and without their “second-search” feature activated, allowing for the identification of multiple peptides from single mixed MS/MS spectra. Furthermore, the novel INFERYS rescoring ^22^ based on fragmentation pattern prediction was evaluated. Lastly, also the performance of SpectroMine was evaluated, but here without employing Percolator for rescoring as SpectroMine runs as a standalone program and not within the PD suite.

## RESULTS & DISCUSSION

### Suppression of Peptide Adsorption to Autosampler Vial Surfaces

In order to ensure minimal sample losses due to adsorption of peptide material to sample vial surfaces and enable reliable processing of protein digest samples at concentrations ≤ 10 ng/μL, we first optimized sample dilution and storage conditions. The positive effects of adding trace amounts of PEG to the sample solvent on sample stability have already been described previously and were evaluated in this study for HeLa tryptic digest samples at a concentration of 250 pg/μL^23^. Aliquots of 100 ng/μL were diluted to 250 pg/μL in sample solvent with 0.001% (w/v) and without PEG and samples were analyzed during a period of 24h after the actual dilution. LC-MS/MS analyses performed at an interval of 4h clearly indicate the positive effect on peptide abundance when using 0.001% PEG as an additive into the sample injection solution (Figure S2). Whereas little or no difference in peptide abundance is observed immediately after sample dilution (abundance ratio PEG/FA close to 1 over the entire elution window), a significant increase in the most hydrophobic portion of the elution window is observed as from 4h after initial sample dilution (abundance ratio PEG/FA increases to 2-3 in the elution window from 35 to 45 minutes). In line with the earlier study on this topic^23^, these results suggest that more hydrophobic peptides are less likely to adsorb to autosampler vial surfaces when trace amounts of PEG are added. This is supported by the observation that the difference in identification numbers grows larger as autosampler residence time is increased (Figure S3). Initial improvements in proteome coverage (0 hrs after sample dilution) observed for the samples containing 0.001% PEG are most likely a result of sample dilution handling. We can however not exclude the possibility that peptide adsorption is affected immediately upon insertion into the sample vial. To quantify the level of contamination introduced by adding PEG to the sample solvent, we have conducted some experiments where blank samples containing 0.001% PEG was injected. When comparing the traces with and without FAIMS, it becomes clear that PEG degradation products elute at regular retention time intervals and that the majority of singly charged PEG ions can be effectively removed by using FAIMS (Figure S4).

### LC Solvent Gradient Optimization Towards Sub Nanogram Proteomics

Next, we evaluated the effect of solvent gradient length to ensure both maximum separation and sufficient concentration to generate clean MS/MS spectra and thereby achieve maximum number of identifications. Even though higher separation resolution is typically achieved by extending the LC solvent gradient, the relative concentration of eluting peptides decreases as a function gradient length ^24–26^. As a result, the concentration of low abundant peptides will drop below the limits of detection at a certain point and further increase in peak capacity will not yield more or even less identifications. The optimal gradient length largely depends on the amount of peptide material injected, which becomes clear when comparing the number of identifications that could be obtained for HeLa digest sample loads of 50, 250 and 500 pg (Figure 1). Even though peak capacity increases from 418 to 638 by extending the gradient length from 30 to 60 minutes (Figure S5), no increase in identifications is observed when 50 pg of HeLa cell digest was injected. However, using longer solvent gradients does have an impact for higher sample loads, resulting in a maximum number of identified protein groups (2449 for 500 pg HeLa cell digest) with a 60 minute solvent gradient. Therefore, the 60 minute gradient LC method was used to explore the proteome coverage that could be achieved for sample loads up to 5 ng of HeLa tryptic peptides. ^1,3,13,27^

### Gas Phase Fractionation Facilitated by FAIMS Pro Improves Sensitivity for Low Sample Loads

In addition to working with LC methods that match the amount of sample that is injected, a significant gain in sensitivity can be achieved by using FAIMS to favor the transmission of multiply charged ions into the MS^28^. This is especially true for limited sample LC-MS/MS analyses, as the relative contribution of singly charged background ions becomes more substantial at low sample loads. This has been demonstrated in figure S5A, where base peak chromatograms for the separation of 1 ng HeLa digest have been compared with and without FAIMS. By applying a single CV (−50 V), the population of ions entering the MS is significantly altered, producing similar peak distribution throughout the peptide elution window, but with a substantial reduction of background chemical noise. This is illustrated by the ‘empty’ appearance of the retention time windows where no peptide elution is expected (Before 10 and after 70 minutes). It can be observed that singly charged background ions (examples indicated in red in the no FAIMS control run in figure S6A) are completely or partially removed without affecting signal intensity to a large extent, resulting in improved signal to noise (S/N) ratios for peptides (illustrated by the increased peptide signal intensity in the region between 10 and 30 minutes). Even though the amount of peptide identifications and PSMs is reduced when analyses are performed at a single CV, protein identifications are typically increased compared to the same analysis without FAIMS (Figure S6B). When injecting 1 ng of HeLa digest, we found that working at a CV of −50 V produces on average 44% more protein group identifications compared to the same analysis without FAIMS. These findings are consistent with earlier reports and confirm that single CV FAIMS reduces the sampling of highly abundant proteins, resulting in higher proteome coverage but with lower peptide sequence coverage per protein^28^. To investigate the potential of using methods with internal CV stepping for the analysis of low sample amounts, a series of exploratory experiments was set up where multiple CVs were used within a single analysis. We evaluated whether scanning between 2 (−50 and −60 V) or 3 (−50, −60 and −70 V) CV values provided greater coverage as compared to a single CV method. When operating the MS in TopN10 DDA mode and injecting 1 ng of HeLa digest, we found that the highest proteome coverage was obtained by using a single CV method at −50 V (Figure S7). To rule out the effect of reduced DDA cycle time per CV, we also briefly investigated the effect of combining internal CV stepping with TopS acquisition mode with fixed MS cycle times (2 and 3s cycle time were evaluated). Neither of the internal stepping CV methods yielded as great a coverage as observed for the single CV method (fixed cycle time data not shown). Even though a significant increase in coverage has been reported by other groups when using internal CV stepping methods^27^, we did not observe this under the conditions tested in the current manuscript.

### Optimized Search Strategies Enhance Proteome Coverage

As different search strategies might also impact the sensitivity achieved in low input experiments, we evaluated various data processing workflows for their ability to identify spectra from low sample amounts. The benchmark results (Figure 2) show a clear benefit for modern database search algorithms and their capability to identify multiple peptides from single mixed fragment spectra (“second search”). Strikingly, this performance improvement increases with concentration (+4.5% peptides at 250 pg to +24% more peptides at 5 ng for Sequest HT) probably due to more frequent occurrence of mixed spectra and wider isolation window for MS2 fragmentation (2 Da). Interestingly, second search leads to decreased performance at 50 pg. An opposite effect is observed for considering predicted fragmentation patterns (INFERYS), where lower concentrations show higher benefit (+22% at 50 pg to +4% at 5 ng, also Sequest HT). At 50 pg, Mascot did not produce enough identifications to allow Percolator separation between targets and decoys and therefore no high confident identifications could be deduced. MS Amanda and SpectroMine can achieve comparable performance at sample loads exceeding 1 ng, but at sample loads below 1 ng, Sequest HT together with INFERYS results in the highest proteome coverage. Consequently, we continued further analysis using this Sequest HT / INFERYS workflow and coupled it to IMP-apQuant ^29^ for label-free quantification and evaluation of chromatographic parameters (i.e. FWHM).

**Figure 2.**
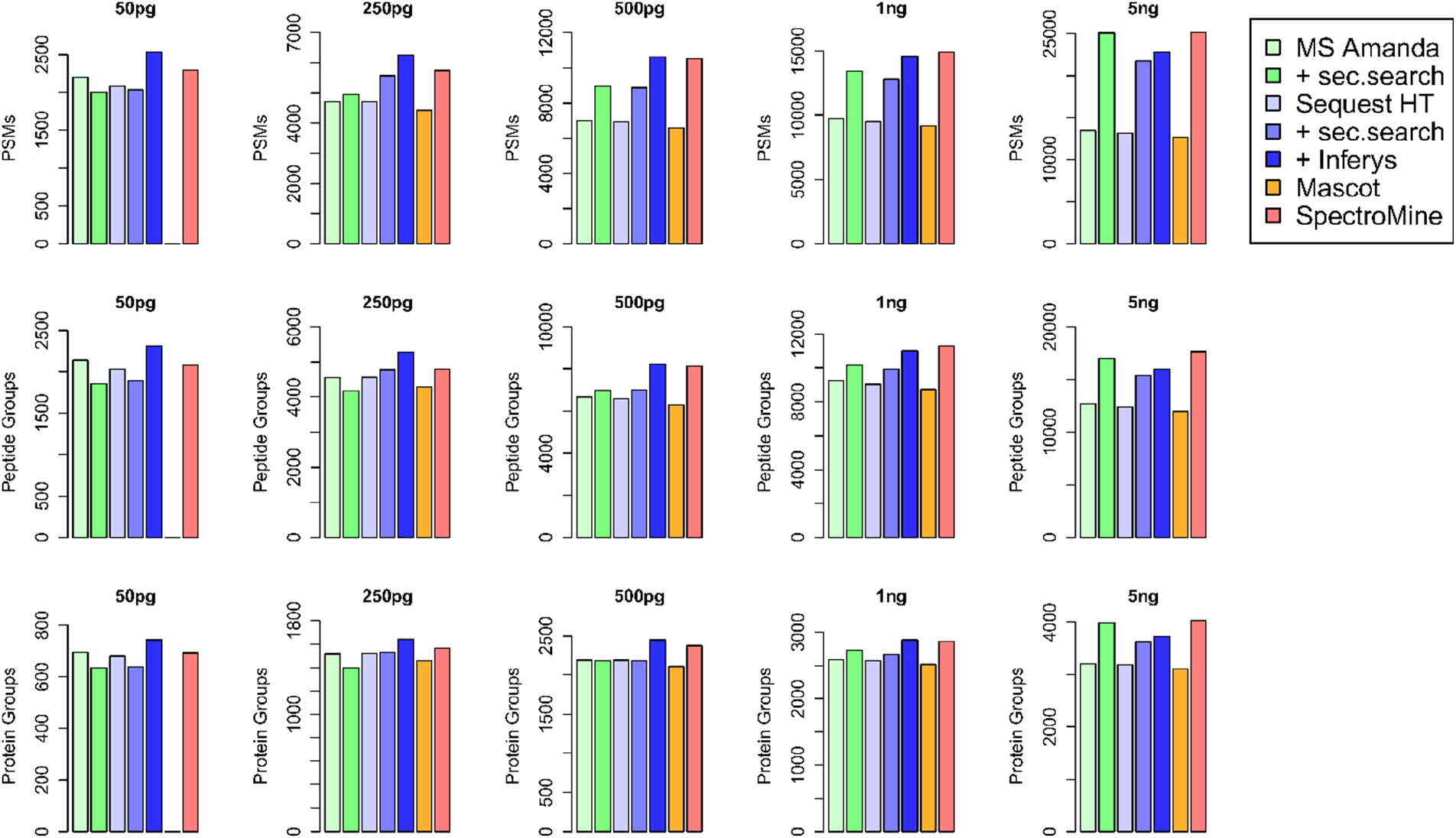
Impact of different search strategies on the sensitivity achieved in low input experiments. MS/MS spectra from raw data acquired at various concentrations of HeLa tryptic digest were analyzed in Proteome Discoverer (PD) to evaluate MS Amanda, Sequest HT, Mascot 2.2 and SpectroMine (not in PD). MS Amanda and Sequest HT were evaluated with and without their “second-search” feature activated, INFERYS rescoring based on fragmentation pattern prediction was evaluated with Sequest HT.

### Deep and Robust Proteome Profiling of Low Sample Amounts

When implementing the Sequest HT / INFERYS workflow for a dilution series of HeLa tryptic digest (0.25, 0.5, 1, 2.5 and 5 ng), we demonstrate identification and quantification of 1486 protein groups on average from just 250 pg (Figure 3a), which approaches the amount of protein expressed in a single cell ^30^. However, one should be aware of the fact that the numbers obtained in current manuscript represent what could be feasible when all pieces of a single cell proteomics workflow have been carefully optimized. Even though several innovative breakthroughs have been described that allow near loss-less single-cell sample preparation^1,3,13,14^, the analysis of sub nanogram aliquots of pre-prepared bulk samples will inevitably be more reproducible and result in a higher yield. This has been illustrated by the results reported in a recent article by Cong et al.^15^, where significant improvements in low input proteome coverage were described by implementing ultra-low flow (ULF) LC (20 nL/min) combined with FAIMS and Orbitrap Eclipse MS. Within a total analysis time of approximately 2.5 hrs, they were able to successfully identify 2061 protein groups using 0.5 ng aliquots of HeLa protein digest. By using a similar gradient length of 1 hr and injecting the same amount of HeLa digest sample (0.5 ng aliquot), we managed to identify 2210 protein groups within a total analysis time of 1.5 hrs. The value of using standardized protein digest samples for benchmarking is evidenced by the observation that the ULF-FAIMS-Eclipse workflow enabled successful identification of 1056 protein groups from a single cell. Therefore, we believe these results can be used as an indication of how protein group identifications arising from pre-prepared bulk samples relate to those that can be obtained for single cell samples.

**Figure 3.**
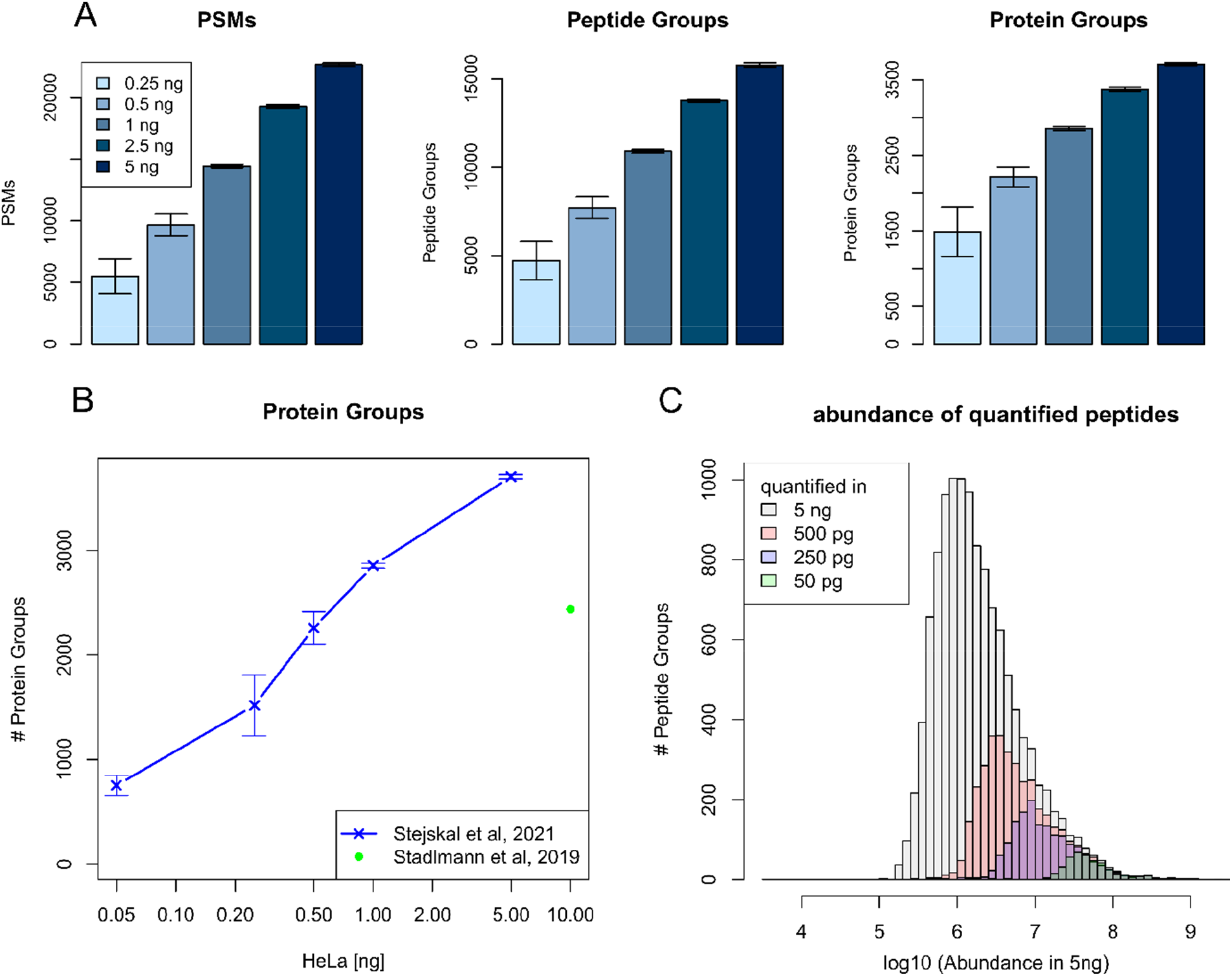
A) PSM, peptide and protein group identifications obtained for a dilution series of HeLa tryptic digest (5 to 0.25 ng injected), Sequest HT with INFERYS rescoring was used as processing workflow in PD 2.5. (average values of technical quadruplicates, n=4) B) The amount of proteins identified as a function of the sample load (ng). Current data is compared to results obtained in 2019 by our group (14). C) Comparison of the abundance of quantified peptides obtained for a 5 ng sample to their detection in lower sample amounts (50, 250, 500 pg).

Overall, the results obtained in the current contribution clearly demonstrate the progress achieved due to the improved chromatographic setup and combination of Orbitrap MS with FAIMS, as now we can identify 2888 protein groups from 1 ng of peptides, whereas in the previous contribution, 2436 protein groups could be identified from 10 ng in a 1 hr gradient ^17^ (Figure 3b). ^28^

It has to be noted that all identification numbers reported in the current manuscript include those arising from a list of common contaminants. We found that the relative contribution of non-human contaminants was marginal but increased with reduced sample load (Figure S9). For the lowest amount (50 pg), 5% of the reported proteins were of non-human origin, mostly from *Bos taurus* and presumably originating from the cell culture medium used to grow the HeLa cells. This number quickly went down below 2% of reported protein groups when sample load was increased to 0.5 ng. To prove that these identifications did not result from system contamination, search results obtained for a blank run immediately after the run where the highest amount of sample was injected (5 ng HeLa tryptic digest) have been included in the supplementary information (Supplementary information figure 10). At this concentration, sample carry-over was found to be 0.47% at the PSM level (107 PSMs), 0.66% at the peptide level (105 peptide groups) and 1.71% at the protein groups level (64 protein groups).

To assess proteome coverage achieved from low input samples, we overlayed peptide abundance in a 5 ng/μL sample with their detection in lower sample amounts (50, 250, 500 pg). Low sample amounts allow for the detection of abundant peptides, whereas less abundant peptide species cannot be detected from low sample amounts (Figure 3c). This is expected and scales with the amount injected. Comparing the ten-percentile apQuant area of peptides detected at low concentration results in 1.3e6 for peptides detected at 500 pg and increases to 1.7e7 for peptides detected in 50 pg corresponding to a factor of 12.6. So, detection of a peptide in a 10-fold lower concentration here requires 12.6-fold higher peptide abundance. It is expected that only abundant peptides can be detected from lower concentrations, especially for homogenous mixtures of cell digest derived from larger cell populations. We hypothesize that single cell samples could enable detection of less abundant peptides, as bulk analysis might mask some higher abundant peptides originating from only a subpopulation of cells. Besides affecting the population of peptides that can be identified, MS1 based quantification reproducibility is also affected when reducing the amount injected. The coefficient of variation obtained for protein quantification in quadruplicate analyses of varying sample loads has been plotted in figure S11. Down to 1 ng of peptide material injected, very reproducible quantification was observed (0.23%CV). When lowering the injected amount below 1 ng, quantification reproducibility decreases steadily to 0.33 and 0.37% for 500 and 250 pg respectively. This is expected and can be attributed to the fact that it is less likely for low abundant peptides to generate clean MS chromatograms that allow proper quantification.

## CONCLUSION

Facilitated by the evolution of LC-MS/MS instrumentation to a level where consistent identification of proteins from sub nanogram protein samples has become feasible, limited sample and single cell proteomics workflows are rapidly emerging from academic and industrial expertise centers. In addition to providing highly specialized sample preparation methods that allow near loss-less and automated processing of cells and proteins, ultra-sensitive separation techniques are mandatory to translate minute sample amounts into qualitative MS signals that can be used for protein identification. Several research groups have demonstrated the potential of using ultra low flow LC as an effective approach to increase detection sensitivity in LC-MS based proteomics. However, LC operation at very low flow rates is far from routine and in many cases still lacks the throughput required for effective processing of large sample batches. An alternative approach using more conventional nanoLC flow rates (250 nL/min) is presented in the current contribution. Bycombiningoptimized sample dilution and storage conditions, a dedicated second generation limited sample micro pillar array column, Orbitrap Exploris™ 480 mass spectrometer, FAIMS Pro interface and the latest database search algorithms, we provide a very promising solution for the analysis of low input protein digest samples. More than a 10-fold increase in detection sensitivity was achieved as compared to a previous contribution, where we reported on identification of 2436 proteins from 10 ng of HeLa tryptic digest. Even though the set-up used was completely different, major improvements in detection sensitivity can be attributed to the use of FAIMS (+40%) and the implementation of a novel type of LC column (+30%). By identifying 1486 and 2210 proteins from as little as 250 and 500 pg of HeLa tryptic digest respectively, we demonstrate the potential of the current setup for integration in limited sample and single cell proteomics workflows.

## SUPPORTING INFORMATION

The Supporting Information is available free of charge at http://pubs.acs.org/

- Design and performance specifications of non-porous pillar array column type
- Effect of PEG addition to sample solvent
- Use of FAIMS at 1CV versus no FAIMS and multiple CV FAIMS
- Coefficient of variance analysis of label-free protein quantification
- Contribution of non-human contaminants to identification number
- System carry-over

## AUTHOR INFORMATION

### Notes

JODB is an employee of PharmaFluidics. PJ is co-founder of PharmaFluidics.

## ACKNOWLEDGEMENTS

We thank Dr. Elisabeth Roitinger, Dr. Markus Hartl and Claudia Ctortecka for proofreading and for fruitful discussions during manuscript preparation. We also thank Julia Kraegenbring from Thermo Fisher for her helping out with setting-up FAIMS Pro and Richard Imre for PRIDE submission. This work was financially supported by the EPIC-XS, project number 823839, the Horizon 2020 Program of the European Union, and the ERA-CAPS I 3686 project of the Austrian Science Fund.

PharmaFluidics acknowledges Vlaio (Flanders Innovation & Entrepreneurship) for the financial support of the development of the second generation pillar array based products (project number HBC.2020.2054).

Spectrometry proteomics data have been deposited to the ProteomeXchange Consortium via the PRIDE ^31^ partner repository with the dataset identifier PXD024017.

## REFERENCES

(1) Brunner, A.; Thielert, M.; Vasilopoulou, C.; Ammar, C.; Coscia, F.; Mund, A.; Horning, O. B.; Bache, N.; Apalategui, A.; Lubeck, M.; Raether, O.; Park, M. A.; Richter, S.; Fischer, D. S.; Theis, F. J.; Meier, F.; Mann, M. Ultra-High Sensitivity Mass Spectrometry Quantifies Single- Cell Proteome Changes upon Perturbation. bioRxiv 2020. https://doi.org/10.1101/2020.12.22.423933.

(2) Bekker-Jensen, D. B.; Martínez-Val, A.; Steigerwald, S.; Rüther, P.; Fort, K. L.; Arrey, T. N.; Harder, A.; Makarov, A.; Olsen, J. v. A Compact Quadrupole-Orbitrap Mass Spectrometer with FAIMS Interface Improves Proteome Coverage in Short LC Gradients. Molecular and Cellular Proteomics 2020, 19 (4), 716–729. https://doi.org/10.1074/mcp.TIR119.001906.

(3) Cong, Y.; Liang, Y.; Motamedchaboki, K.; Huguet, R.; Zhao, R.; Shen, Y.; Lopez-ferrer, D.; Zhu, Y.; Kelly, R. T. Improved Single Cell Proteome Coverage Using Narrow-Bore Packed NanoLC Columns and Ultrasensitive Mass Spectrometry. 2020. https://doi.org/10.1021/acs.analchem.9b04631.

(4) Dou, M.; Clair, G.; Tsai, C. F.; Xu, K.; Chrisler, W. B.; Sontag, R. L.; Zhao, R.; Moore, R. J.; Liu, T.; Pasa-Tolic, L.; Smith, R. D.; Shi, T.; Adkins, J. N.; Qian, W. J.; Kelly, R. T.; Ansong, C.; Zhu, Y. High-Throughput Single Cell Proteomics Enabled by Multiplex Isobaric Labeling in a Nanodroplet Sample Preparation Platform. Analytical Chemistry 2019, 91 (20), 13119–13127. https://doi.org/10.1021/acs.analchem.9b03349.

(5) Zhu, Y.; Zhao, R.; Piehowski, P. D.; Moore, R. J.; Lim, S.; Orphan, V. J.; Paša-Tolić, L.; Qian, W. J.; Smith, R. D.; Kelly, R. T. Subnanogram Proteomics: Impact of LC Column Selection, MS Instrumentation and Data Analysis Strategy on Proteome Coverage for Trace Samples. International Journal of Mass Spectrometry 2018, 427 (February), 4–10. https://doi.org/10.1016/j.ijms.2017.08.016.

(6) Aebersold, R.; Mann, M. Mass-Spectrometric Exploration of Proteome Structure and Function. Nature 2016, 537 (7620), 347–355. https://doi.org/10.1038/nature19949.

(7) Angel, T. E.; Aryal, U. K.; Hengel, S. M.; Baker, E. S.; Kelly, R. T.; Robinson, E. W.; Smith, R. D. Mass Spectrometry-Based Proteomics: Existing Capabilities and Future Directions. Chemical Society Reviews 2012, 41 (10), 3912–3928. https://doi.org/10.1039/c2cs15331a.

(8) Dalerba, P.; Kalisky, T.; Sahoo, D.; Rajendran, P. S.; Rothenberg, M. E.; Leyrat, A. A.; Sim, S.; Okamoto, J.; Johnston, D. M.; Qian, D.; Zabala, M.; Bueno, J.; Neff, N. F.; Wang, J.; Shelton, A. A.; Visser, B.; Hisamori, S.; Shimono, Y.; Van De Wetering, M.; Clevers, H.; Clarke, M. F.; Quake, S. R. Single-Cell Dissection of Transcriptional Heterogeneity in Human Colon Tumors. Nature Biotechnology 2011, 29 (12), 1120–1127. https://doi.org/10.1038/nbt.2038.

(9) Goebel-Stengel, M.; Stengel, A.; Taché, Y.; Reeve, J. R. The Importance of Using the Optimal Plasticware and Glassware in Studies Involving Peptides. Analytical Biochemistry 2011, 414 (1), 38–46. https://doi.org/10.1016/j.ab.2011.02.009.

(10) Zhu, Y.; Piehowski, P. D.; Zhao, R.; Chen, J.; Shen, Y.; Moore, R. J.; Shukla, A. K.; Petyuk, V. A.; Campbell-Thompson, M.; Mathews, C. E.; Smith, R. D.; Qian, W. J.; Kelly, R. T. Nanodroplet Processing Platform for Deep and Quantitative Proteome Profiling of 10-100 Mammalian Cells. Nature Communications 2018, 9 (1), 1–10. https://doi.org/10.1038/s41467-018-03367-w.

(11) Budnik, B.; Levy, E.; Harmange, G.; Slavov, N. SCoPE-MS: Mass Spectrometry of Single Mammalian Cells Quantifies Proteome Heterogeneity during Cell Differentiation 06 Biological Sciences 0601 Biochemistry and Cell Biology 06 Biological Sciences 0604 Genetics. Genome Biology 2018, 19 (1), 1–12. https://doi.org/10.1186/s13059-018-1547-5.

(12) Specht, H.; Emmott, E.; Petelski, A. A.; Gray Huffman, R.; Perlman, D. H.; Serra, M.; Kharchenko, P.; Koller, A.; Slavov, N. Single-Cell Mass-Spectrometry Quantifies the Emergence of Macrophage Heterogeneity. bioRxiv 2019. https://doi.org/10.1101/665307.

(13) Liang, Y.; Acor, H.; McCown, M. A.; Nwosu, A. J.; Boekweg, H.; Axtell, N. B.; Truong, T.; Cong, Y.; Payne, S. H.; Kelly, R. T. Fully Automated Sample Processing and Analysis Workflow for Low-Input Proteome Profiling. Analytical Chemistry 2021, 93 (3), 1658–1666. https://doi.org/10.1021/acs.analchem.0c04240.

(14) Hartlmayr, D.; Ctortecka, C.; Seth, A.; Mendjan, S.; Tourniaire, G.; Mechtler, K.; Biocenter, V. An Automated Workflow for Label-Free and Multiplexed Single Cell Proteomics Sample Preparation at Unprecedented Sensitivity. bioRxiv 2021, 2021.04.14.439828.

(15) Cong, Y.; Motamedchaboki, K.; Misal, S. A.; Liang, Y.; Guise, A. J.; Truong, T.; Huguet, R.; Plowey, E. D.; Zhu, Y.; Lopez-Ferrer, D.; Kelly, R. T. Ultrasensitive Single-Cell Proteomics Workflow Identifies >1000 Protein Groups per Mammalian Cell. Chemical Science 2021, 12 (3), 1001–1006. https://doi.org/10.1039/d0sc03636f.

(16) Wilm, M.; Mann, M. Analytical Properties of the Nanoelectrospray Ion Source. Analytical Chemistry 1996, 68 (1), 1–8. https://doi.org/10.1021/ac9509519.

(17) Stadlmann, J.; Hudecz, O.; Krššáková, G.; Ctortecka, C.; Van, G.; Beeck, J. op de; Desmet, G.; Penninger, J. M.; Jacobs, P.; Mechtler, K. Improved Sensitivity in Low-Input Proteomics Running Title : Improved Sensitivity in Low-Input Proteomics Using Micro-Pillar Array-Based Affiliations : Analytical Biochemistry 2019, 91 (22), 14203–14207. https://doi.org/10.1021/acs.analchem.9b02899.

(18) MacCoss M, M. J.; Noble, W. S.; Käll, L. Fast and Accurate Protein False Discovery Rates on Large-Scale Proteomics Data Sets with Percolator 3.0. Journal of the American Society for Mass Spectrometry 2016, 27 (11), 1719–1727. https://doi.org/10.1007/s13361-016-1460-7.

(19) Dorfer, V.; Pichler, P.; Stranzl, T.; Stadlmann, J.; Taus, T.; Winkler, S.; Mechtler, K. MS Amanda, a Universal Identification Algorithm Optimized for High Accuracy Tandem Mass Spectra. Journal of Proteome Research 2014, 13 (8), 3679–3684. https://doi.org/10.1021/pr500202e.

(20) Eng, J. K.; McCormack, A. L.; Yates, J. R. An Approach to Correlate Tandem Mass Spectral Data of Peptides with Amino Acid Sequences in a Protein Database. Journal of the American Society for Mass Spectrometry 1994, 5 (11), 976–989. https://doi.org/10.1016/1044-0305(94)80016-2.

(21) Koenig, T.; Menze, B. H.; Kirchner, M.; Monigatti, F.; Parker, K. C.; Patterson, T.; Steen, J. J.; Hamprecht, F. A.; Steen, H. Robust Prediction of the MASCOT Score for an Improved Quality Assessment in Mass Spectrometric Proteomics. Journal of Proteome Research 2008, 7 (9), 3708–3717. https://doi.org/10.1021/pr700859x.

(22) Gessulat, S.; Schmidt, T.; Zolg, D. P.; Samaras, P.; Schnatbaum, K.; Zerweck, J.; Knaute, T.; Rechenberger, J.; Delanghe, B.; Huhmer, A.; Reimer, U.; Ehrlich, H.; Aiche, S.; Kuster, B.; Wilhelm, M. Prosit: Proteome-Wide Prediction of Peptide Tandem Mass Spectra by Deep Learningt. Nature Methods 2019, 16 (6), 509–518. https://doi.org/10.1038/s41592-019-0426-7.

(23) Stejskal, K.; Potěšil, D.; Zdráhal, Z. Suppression of Peptide Sample Losses in Autosampler Vials. Journal of Proteome Research 2013, 12 (6), 3057–3062. https://doi.org/10.1021/pr400183v.

(24) Neue, U. D. Theory of Peak Capacity in Gradient Elution. Journal of Chromatography A 2005, 1079 (1-2 SPEC. ISS.), 153–161. https://doi.org/10.1016/j.chroma.2005.03.008.

(25) Neue, U. D. Peak Capacity in Unidimensional Chromatography. Journal of Chromatography A 2008, 1184 (1–2), 107–130. https://doi.org/10.1016/j.chroma.2007.11.113.

(26) Petersson, P.; Frank, A.; Heaton, J.; Euerby, M. R. Maximizing Peak Capacity and Separation Speed in Liquid Chromatography. Journal of Separation Science 2008, 31 (13), 2346–2357. https://doi.org/10.1002/jssc.200800064.

(27) Greguš, M.; Kostas, J. C.; Ray, S.; Abbatiello, S. E.; Ivanov, A. R. Improved Sensitivity of Ultralow Flow LC-MS-Based Proteomic Profiling of Limited Samples Using Monolithic Capillary Columns and FAIMS Technology. Analytical Chemistry 2020, 92 (21), 14702–14712. https://doi.org/10.1021/acs.analchem.0c03262.

(28) Hebert, A. S.; Prasad, S.; Belford, M. W.; Bailey, D. J.; McAlister, G. C.; Abbatiello, S. E.; Huguet, R.; Wouters, E. R.; Dunyach, J. J.; Brademan, D. R.; Westphall, M. S.; Coon, J. J. Comprehensive Single-Shot Proteomics with FAIMS on a Hybrid Orbitrap Mass Spectrometer. Analytical Chemistry 2018, 90 (15), 9529–9537. https://doi.org/10.1021/acs.analchem.8b02233.

(29) Doblmann, J.; Dusberger, F.; Imre, R.; Hudecz, O.; Stanek, F.; Mechtler, K.; Dürnberger, G. ApQuant: Accurate Label-Free Quantification by Quality Filtering. Journal of Proteome Research 2019, 18 (1), 535–541. https://doi.org/10.1021/acs.jproteome.8b00113.

(30) Zhu, Y.; Piehowski, P. D.; Kelly, R. T.; Qian, W. J. Nanoproteomics Comes of Age. Expert Review of Proteomics 2018, 15 (11), 865–871. https://doi.org/10.1080/14789450.2018.1537787.

(31) Perez-Riverol, Y.; Csordas, A.; Bai, J.; Bernal-Llinares, M.; Hewapathirana, S.; Kundu, D. J.; Inuganti, A.; Griss, J.; Mayer, G.; Eisenacher, M.; Pérez, E.; Uszkoreit, J.; Pfeuffer, J.; Sachsenberg, T.; Yilmaz, Ş.; Tiwary, S.; Cox, J.; Audain, E.; Walzer, M.; Jarnuczak, A. F.; Ternent, T.; Brazma, A.; Vizcaíno, J. A. The PRIDE Database and Related Tools and Resources in 2019: Improving Support for Quantification Data. Nucleic Acids Research 2019, 47 (D1), D442–D450. https://doi.org/10.1093/nar/gky1106.

