## Supporting information for "Ultra-sensitive nanoLC-MS of sub nanogram protein samples using second generation micro pillar array LC technology with Orbitrap Exploris 480 and FAIMS PRO"

^2^ PharmaFluidics, Technologiepark-Zwijnaarde 82, B-9052 Gent, Belgium.

^3^ IMP - Institute of Molecular Pathology, Campus-Vienna-Biocenter 1, A-1030 Vienna, Austria.

^4^ Gregor Mendel Institute of Molecular Plant Biology of the Austrian Academy of Sciences, Dr. Bohr Gasse 3, A-1030 Vienna, Austria.

^§^ These authors contributed equally.

**CONTENTS**

**Figure S1………………………………………………………………………………………………S2**

**Figure S2………………………………………………………………………………………………S3**

**Figure S3………………………………………………………………………………………………S4**

**Figure S4………………………………………………………………………………………………S5**

**Figure S5………………………………………………………………………………………………S6**

**Figure S6………………………………………………………………………………………………S7**

**Figure S7………………………………………………………………………………………………S8**

**Figure S8………………………………………………………………………………………………S9**

**Figure S9……………………………………………………………………………………………..S10**

**Figure S10……………………………………………………………………………………………S11**

**Figure S11……………………………………………………………………………………………S12**

**Figure S12……………………………………………………………………………………………S13**

**Limited Sample Micro Pillar Array Column…………………………………………………….S14**

**Chromatographic Metrics ………………………………………………………………………...S14**

**References……………………………………………………………………………………………S15**

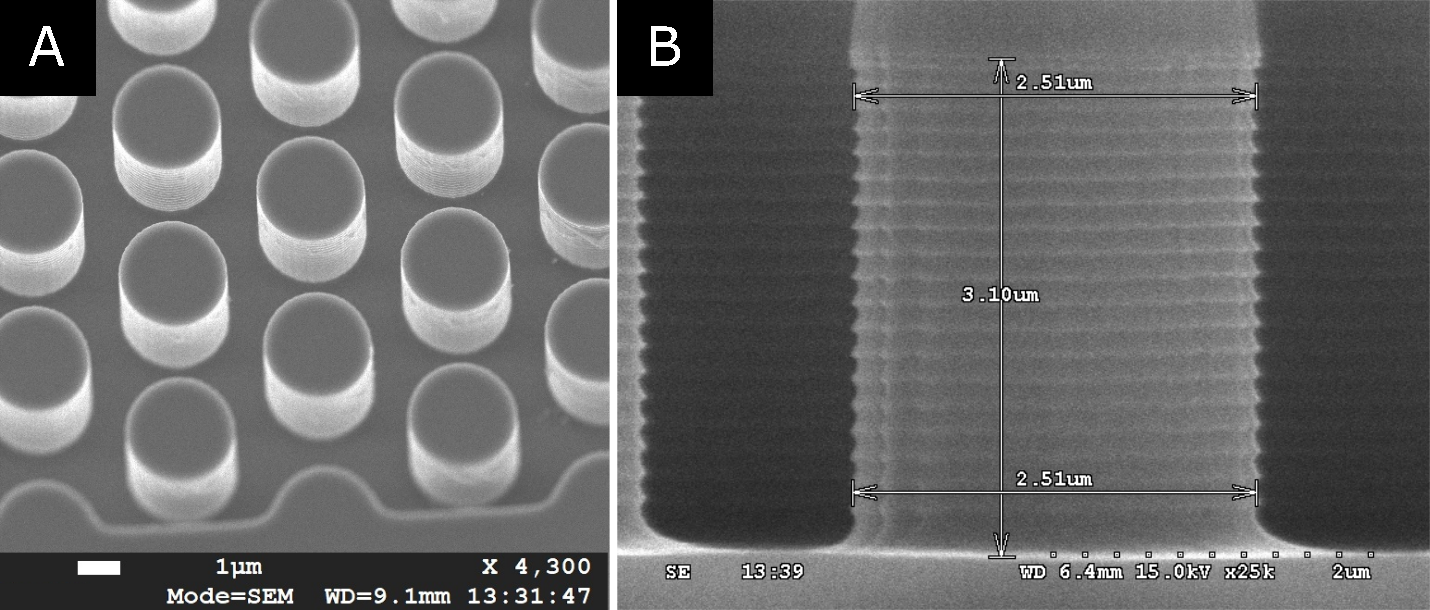

### **Figure S1. A) Scanning Electron Microscopy (SEM) image showing a birds eye view of the second generation micro pillar array stationary phase. B) SEM image showing a transverse section of the micro pillar array column, zoom in on a single 2.5 µm diameter non-porous micro pillar.**

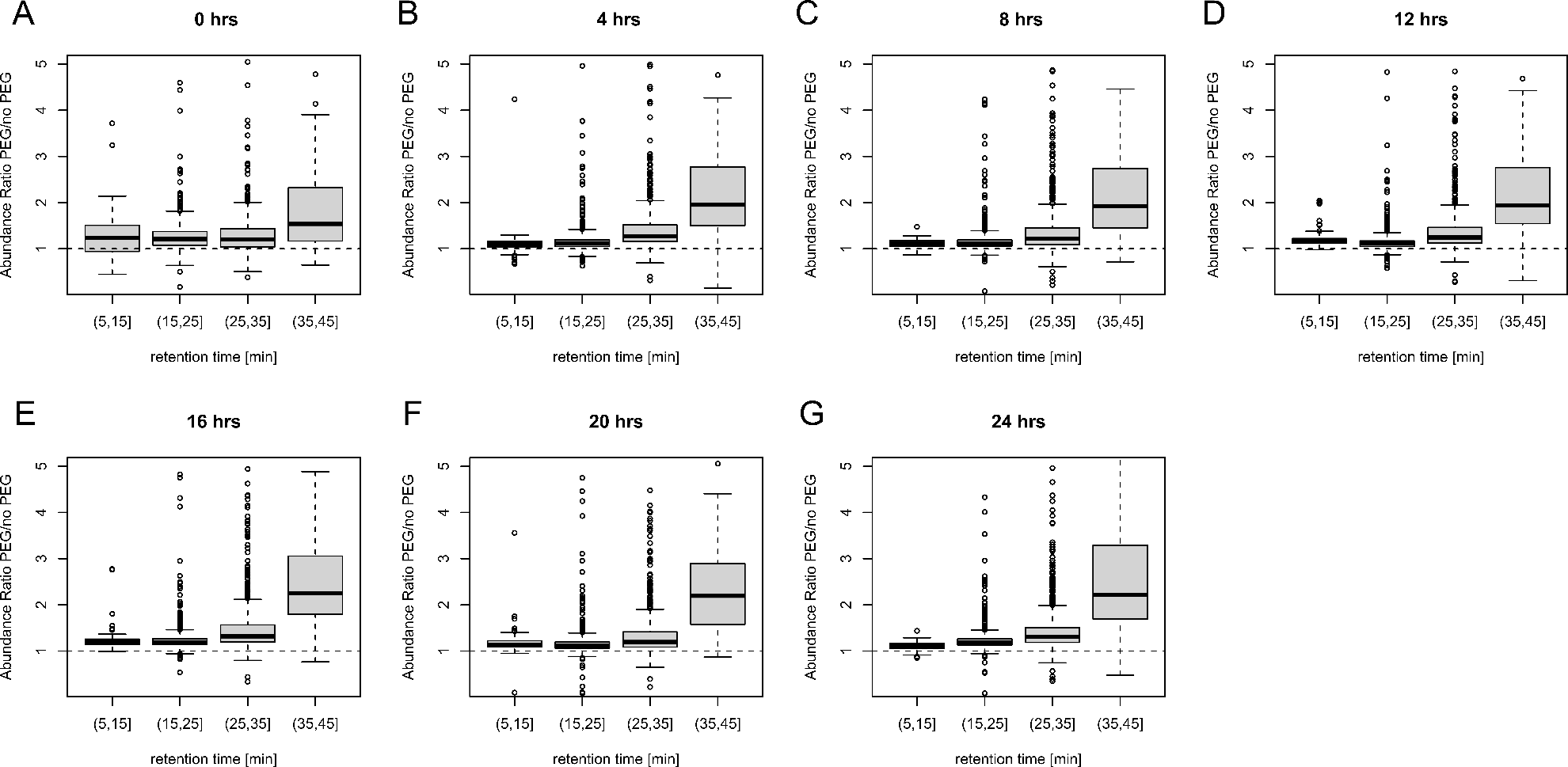

**Figure S2. A-G) Abundance ratio obtained on the peptide level for samples where PEG (0.001% w/v) and no PEG was added to the sample solvent (0.1% FA). Abundance ratio has been plotted for 4 distinct elution windows (5-15,15-25, 25-35 and 35-45 min) representative for the increasing hydrophobicity of peptides. Results obtained over a period of 24h after initial sample dilution are shown at an interval of 4h.**

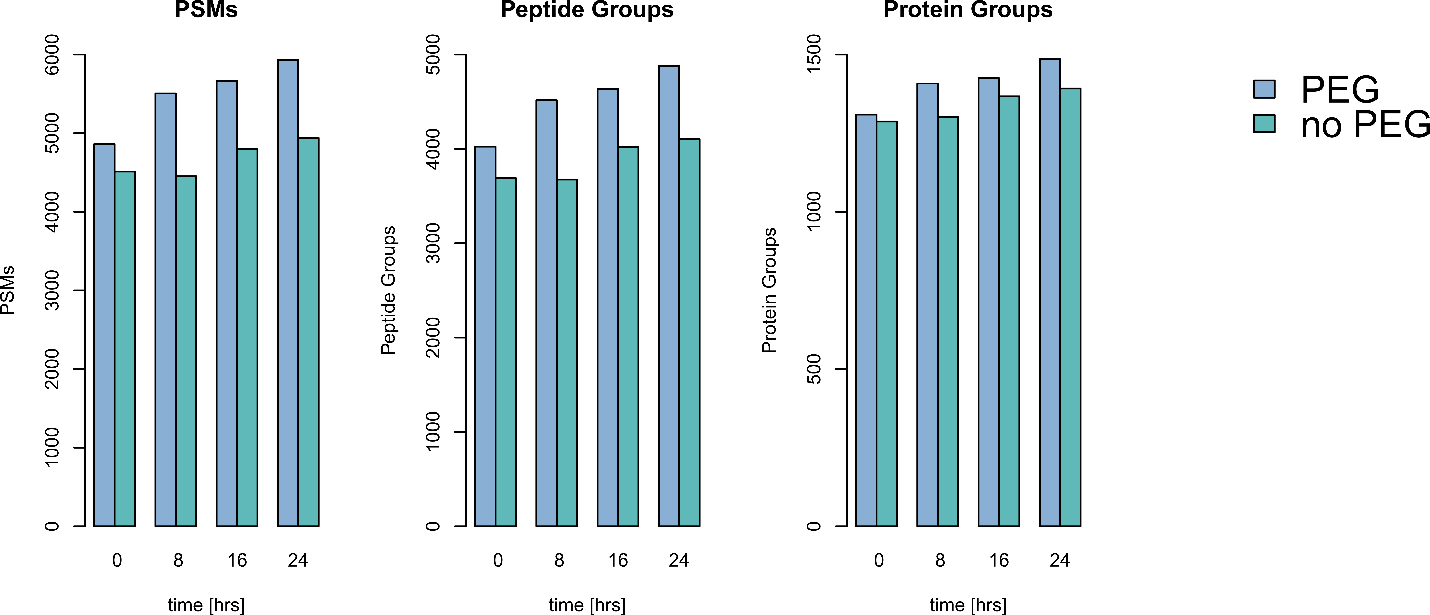

**Figure S3. Evolution of the identification numbers (PSM, peptide and protein groups) obtained for 250 ng of HeLa tryptic digest as a function of time after dilution. Samples where PEG (0.001% w/v) and no PEG was added to the sample solvent (0.1% FA) have been compared.**

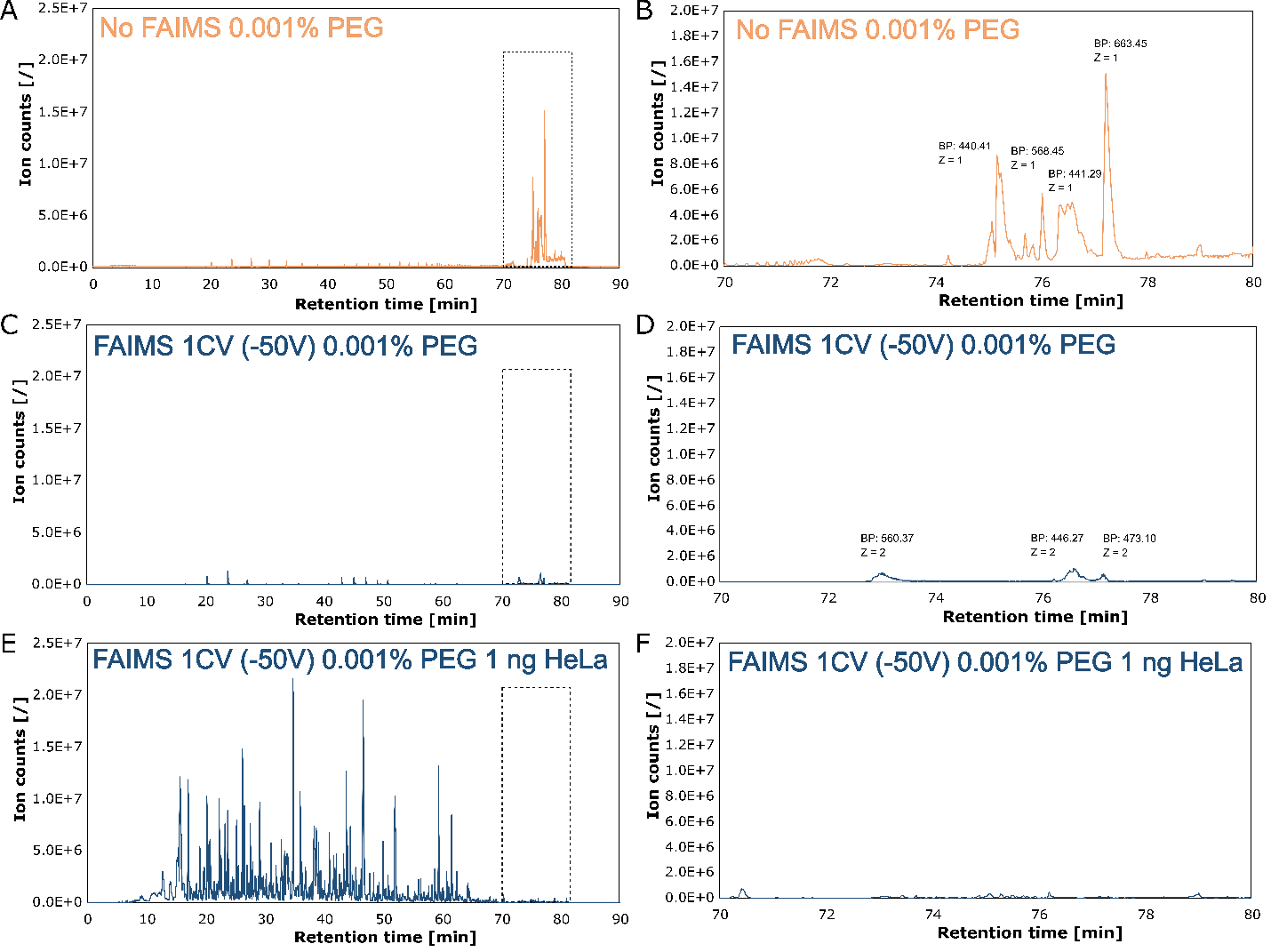

**Figure S4. MS base peak chromatograms obtained for 60 min gradient separation 0.001% PEG in sample solvent (0.1% FA) without FAIMS (A), 0.001% PEG in sample solvent (0.1% FA) with FAIMS 1 CV, -50 V (C) and 1 ng of HeLa tryptic digest, 0.001% PEG in sample solvent (0.1% FA) with FAIMS 1 CV, -50 V. Zoom-in on the high organic wash region (78% Acetonitrile, 0.1% FA) shown to the right (B, D & F).**

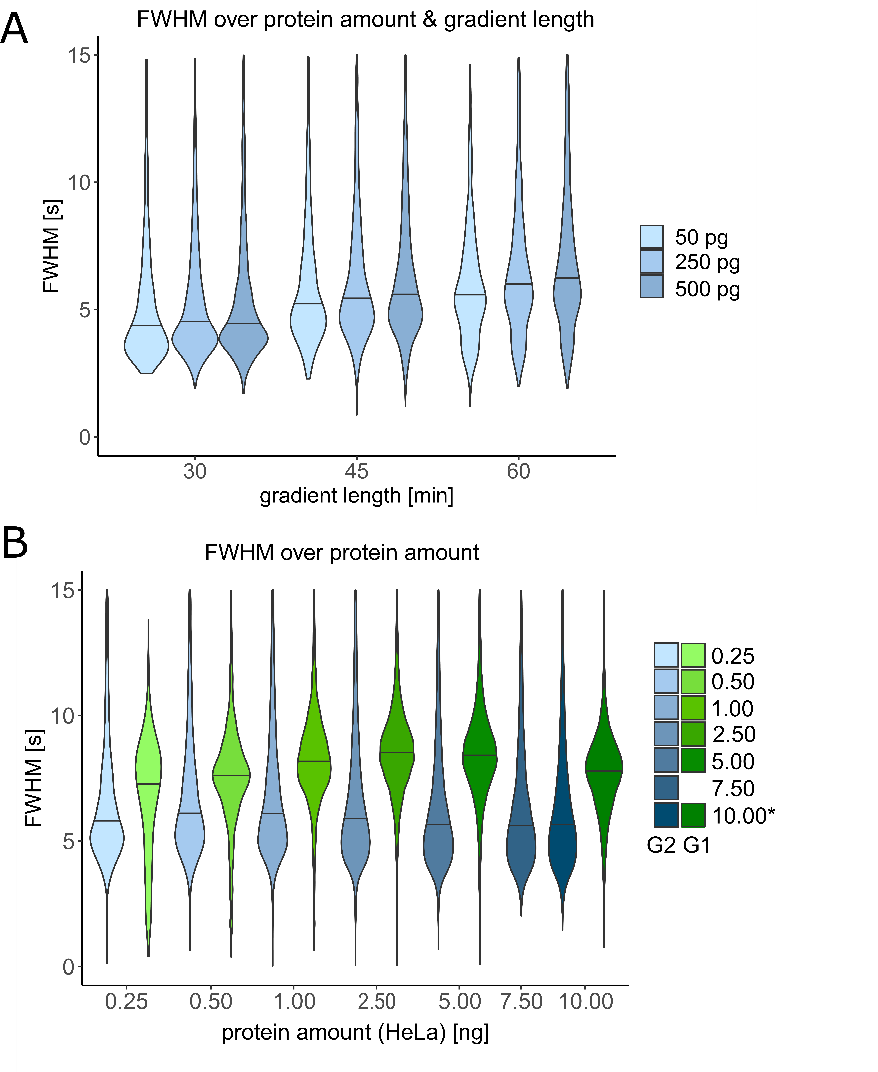

**Figure S5. A) FWHM values obtained for all identified peptides as a function of the sample load (50, 250 and 500 pg HeLa tryptic digest) and gradient length (30, 45 and 60 min). B) FWHM values obtained for all identified peptides as a function of the sample load (0.25, 0.5, 1, 2.5, 5, 7.5, 10 ng HeLa tryptic digest). Data obtained on the dedicated limited sample pillar array column (G2-blue) is compared to results obtained with a first generation superficially porous micro pillar array column (G1-green). *10 ng HeLa tryptic digest data from a previous contribution**^1^ **has also been plotted.**

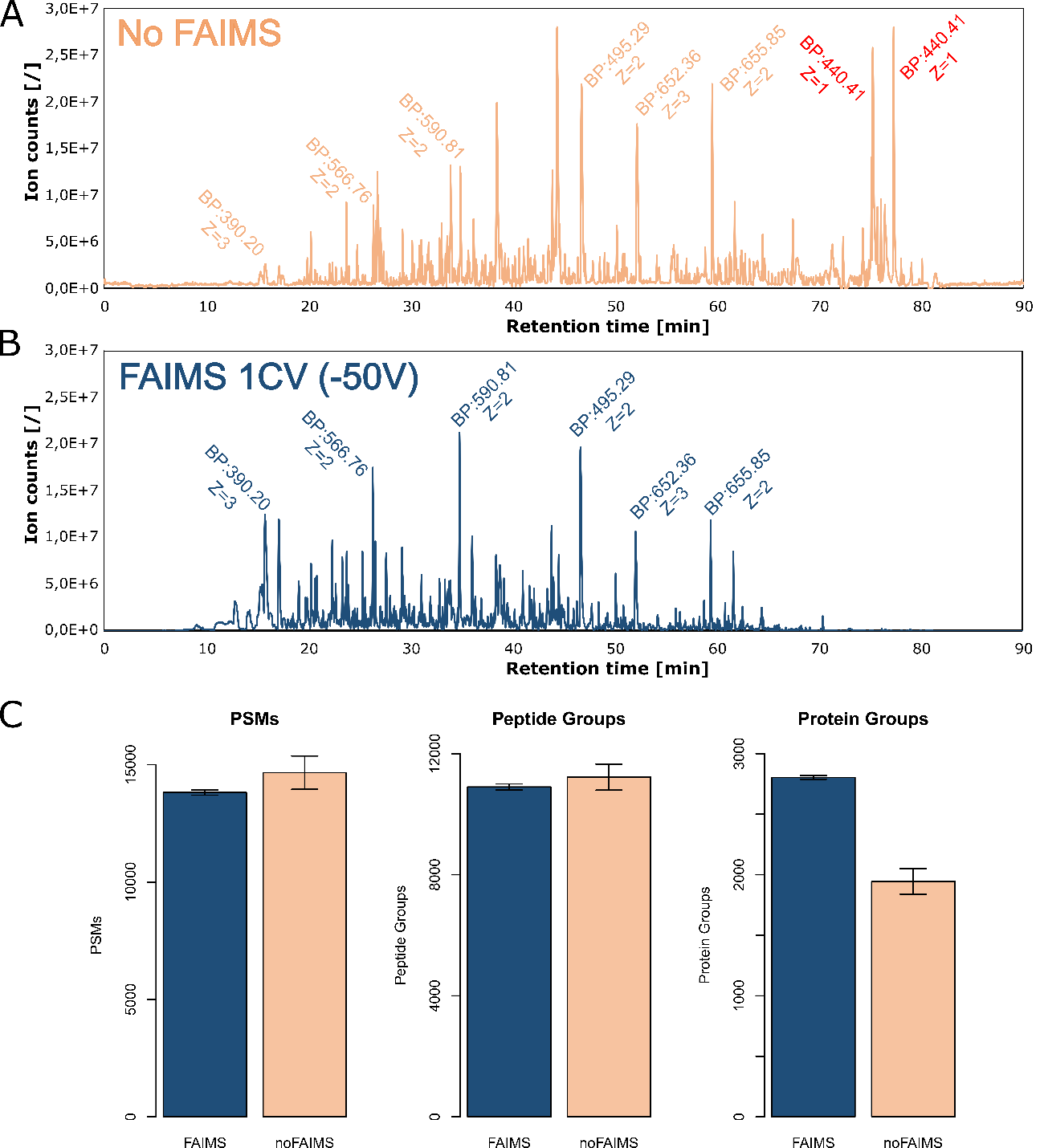

**Figure S6. MS base peak chromatograms obtained for 60 min gradient separation of 1 ng of HeLa tryptic digest, 0.001% PEG in sample solvent (0.1% FA) without (A) and with FAIMS (B). Respective PSM, peptide and protein group identifications have been compared in panel C, With FAIMS: blue, without FAIMS orange.**

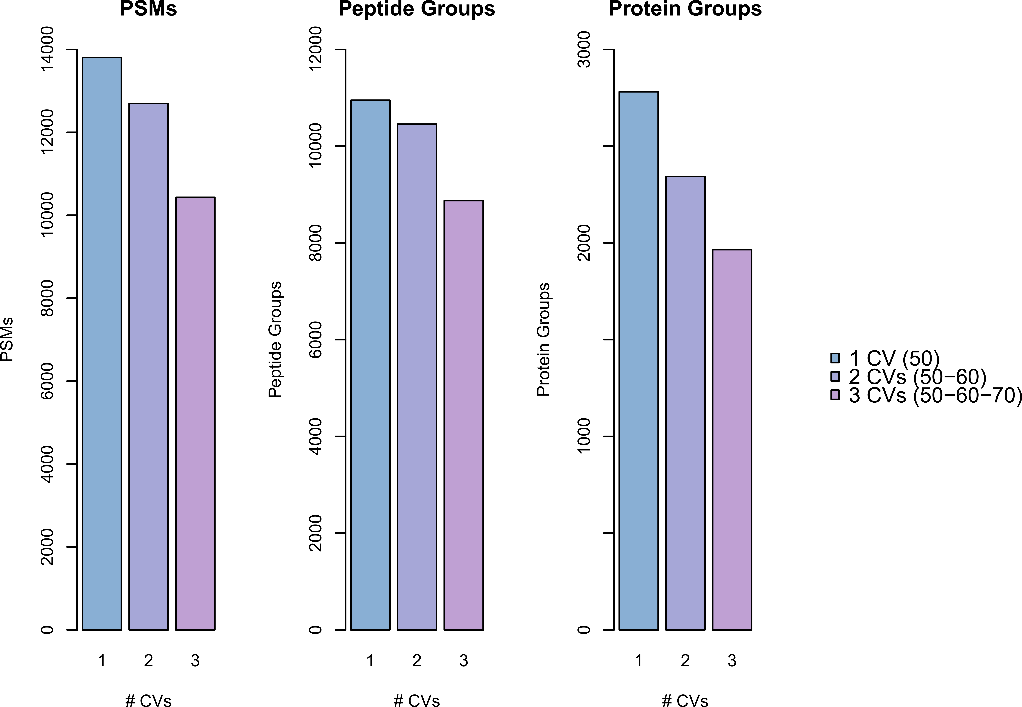

**Figure S7. PSM, peptide and protein group identifications obtained for 1 ng of HeLa tryptic digest using the FAIMS pro interface at either 1 CV (-50V - blue) or 2 CVs (-50 and -60V-purple) and 3 CVs (-50, -60 and -70V-pink) with fast internal CV stepping.**

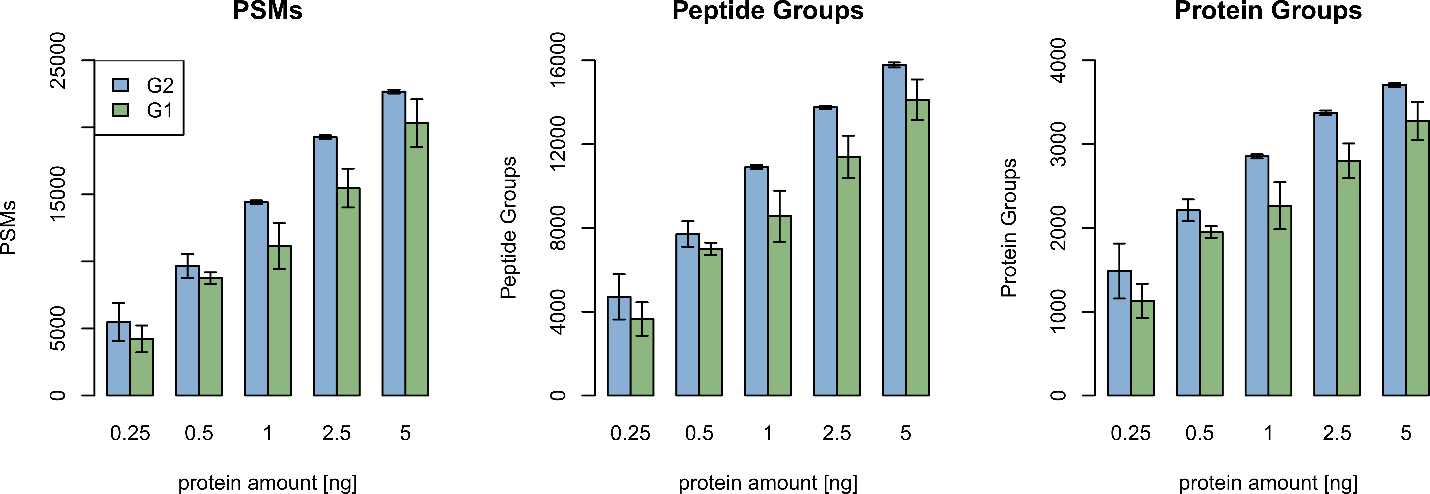

**Figure S8. PSM, peptide and protein group identifications obtained for different amounts of HeLa tryptic digest on either a dedicated limited sample pillar array column (G2-blue: 50 cm length – 2.5 µm pillars – non-porous and a first generation pillar array column (G1-green: 50 cm length – 5 µm pillars – superficially porous) .**

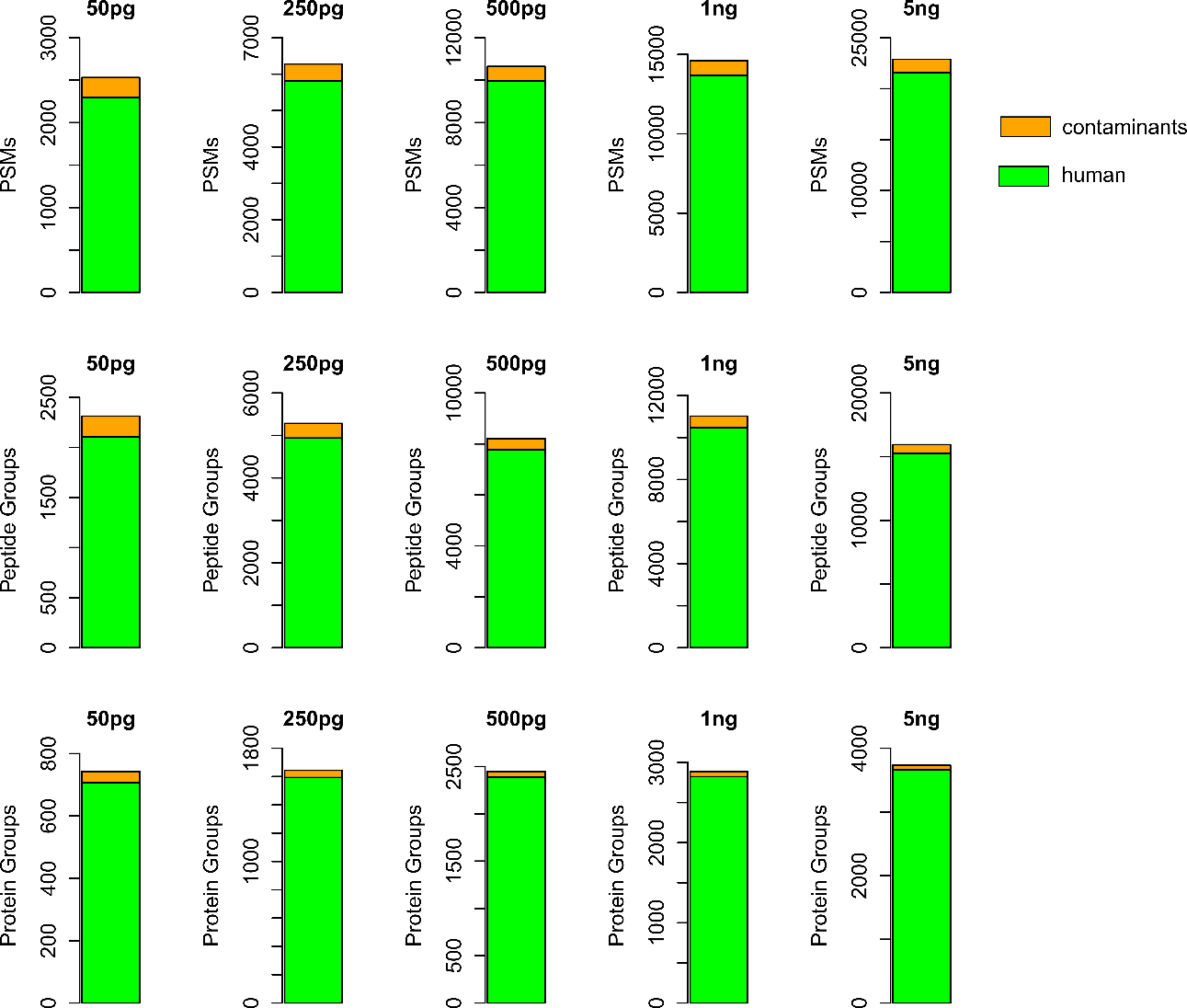

**Figure S9. Relative contribution of non-human contaminants to the overall PSM, peptide and protein group identifications obtained for different amounts of HeLa tryptic digest.**

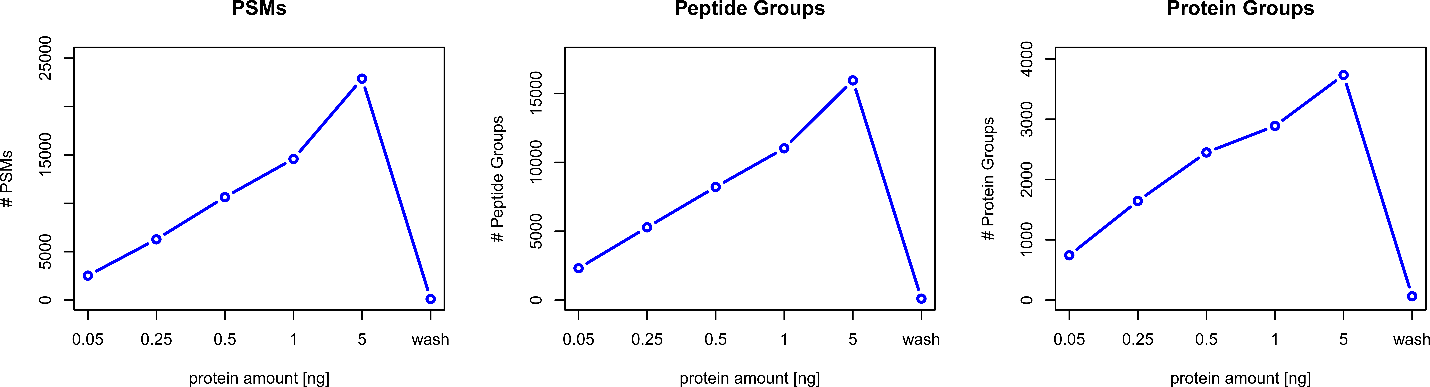

**Figure S10. PSM, peptide and protein groups identifications obtained for different amounts of HeLa tryptic digest and a blank or wash run that was performed immediately after injecting 5 ng of HeLa tryptic digest, Sequest HT with INFERYS rescoring was used as processing workflow in PD 2.5. (average values of technical quadruplicates, n=4)**

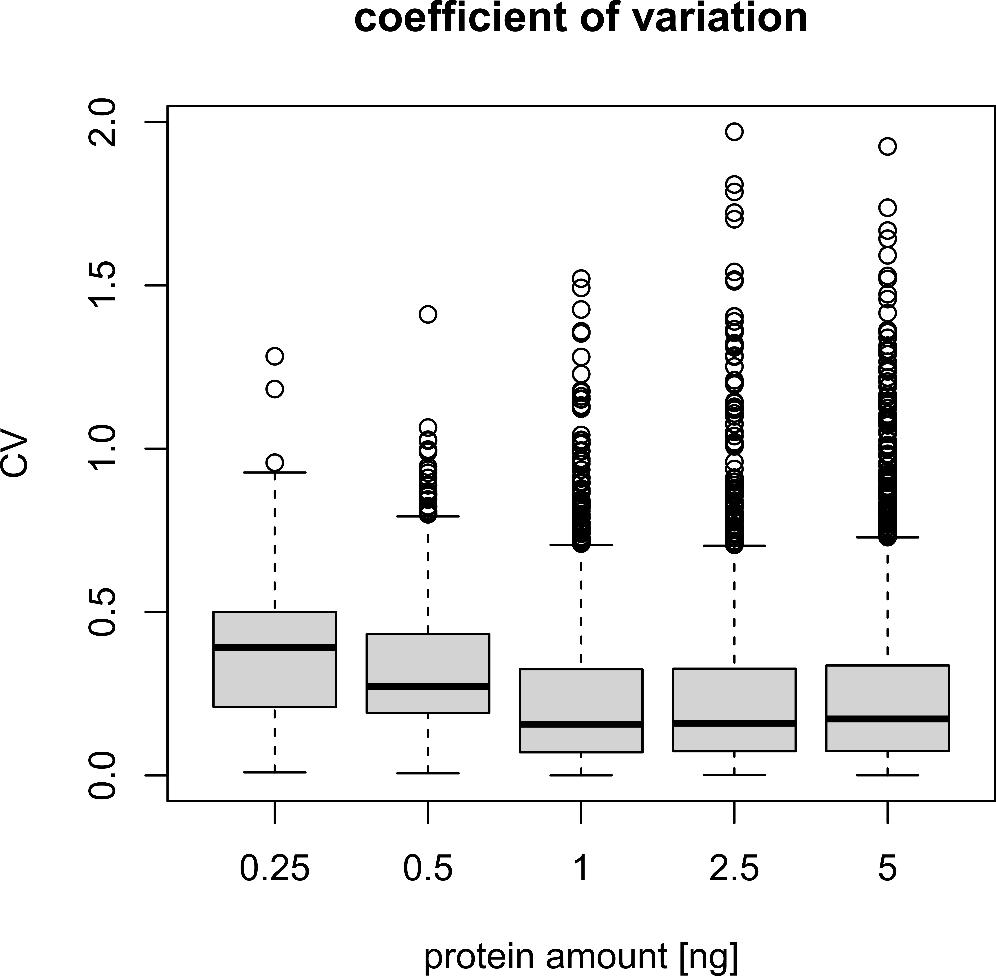

**Figure S11. Coefficient of variation (CV) analysis of MS1 quantitation values across quadruplicates for different amounts of HeLa tryptic digest.**

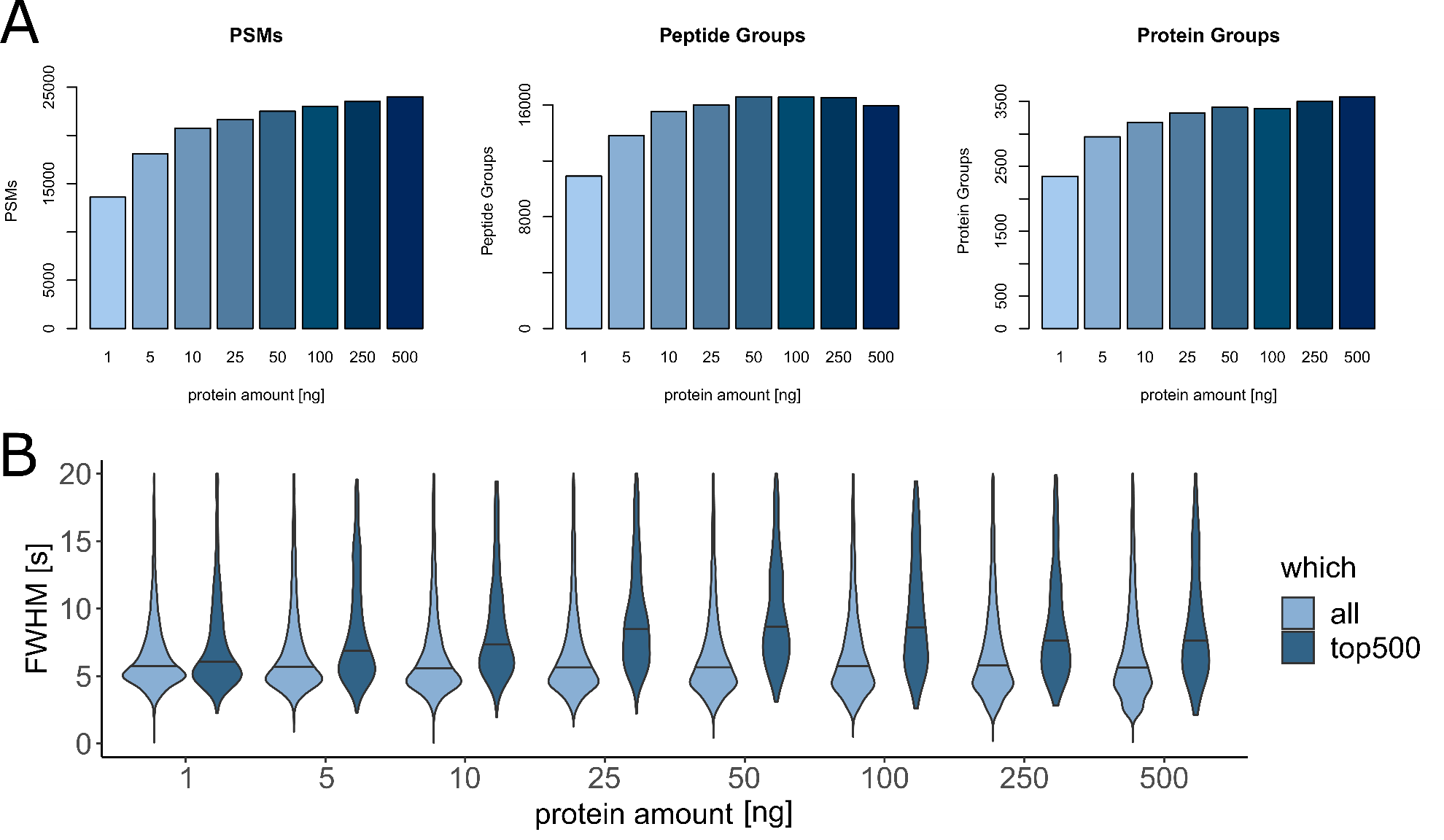

**Figure S12. Evaluation of column loadability. A) PSM, peptide and protein groups identifications obtained for peptides amount ranging from 1 up to 500 ng. TopS instead of TopN10 acquisition method used. B) Evolution of FWHM peptide peak width as a function of the amount of sample injected. FWHM distribution for all peptides (light blue) and top 500 highest abundant peptides (dark blue) are compared.**

**Limited Sample Micro Pillar Array Column**

The nature of the pillar array fabrication process allows precise control over the stationary phase morphology and can therefore be finetuned towards certain applications ^2,3^. In an attempt to further tune the pillar array based stationary phase towards optimal compatibility with very low sample amounts, three important adaptations have been made to the column design. Advanced deep UV lithography has been used to reduce the pillar dimensions by a factor of 2, yielding an array of 2.5 µm diameter silicon pillars positioned at a distance of 1.25 µm (Figure S1). By reducing the distance that molecules have to travel to interact with the stationary phase, the effect of resistance to mass transfer on peak dispersion is reduced is and more efficient chromatography can be achieved ^4^. This ultimately results in an increased signal to noise ratio when entering the MS. By using non-porous instead of superficially porous pillars, an additional gain in sensitivity can be achieved through a complementary effect on peak dispersion. Interaction with the stationary phase is restricted to the outer pillar shell instead of the mesoporous mantle (or the entire mesoporous matrix in the case of fully porous particles). This has a beneficial effect on separation performance separation as long as the interaction surface of the pillars is not fully saturated (and sample overloading starts having a detrimental effect on separation performance). And finally, further reduction of the channel cross section by a factor of 2 ensures optimal peptide elution at flow rates between 100 and 250 nL/min, exploiting the increased electrospray ionization efficiencies associated with low end nanoflow LC ^5^.

**Chromatographic Metrics**

Integration of the IMP-apQuant node ^6^ into the PD processing workflow enables convenient and automated evaluation of chromatographic parameters (i.e. FWHM, kurtosis, skewness) for bottom-up MS/MS analyses of complex protein digest samples. The FWHM distribution obtained for different sample loads (50, 250 and 500 ng HeLa tryptic digest) and different gradient lengths have been plotted in figure S5A and gives a clear representation of the chromatographic performance that can be achieved with the novel micro pillar array column. Based on the median FWHM value that was observed for a 30, 45 and 60 min solvent gradient, peak capacity values ranging from 400 for the shortest to 640 for the longest gradient could be obtained. When comparing these data to the results we previously published using a micro pillar array column with superficially porous pillars and pillar dimensions that are twice the size of those of the current column, a significant reduction of the FWHM values (from 7.94 to 5.64 s; 29% reduction) is be observed. These values are not in full agreement with what was published in 2019, because improved FWHM calculation in apQuant is used. Whereas the previous version only measured time between data points that are above half maximum, the current version uses linear interpolation on both sides which is more accurate. A net gain in peak capacity of 41% (from 543 to 638) is demonstrated, which can mainly be attributed to the reduction of the pillar diameter and interpillar spacing (Figure S5B). Further downscaling of the channel cross section and the absence of a meso porous layer has an effect on the mass loadability of the column (reduction by a factor of approximately 30). Excellent chromatography with no signs of excessive column overloading can however be achieved for samples with a total protein content up to 10 ng. This is illustrated in figure S5B, where FWHM distributions obtained for a HeLa tryptic digest dilution series from 10 down to 0.25 ng are compared. To determine the maximum loading capacity of the column, and additional experiment was performed where the amount of HeLa digest injected was systematically increased up to 500 ng. Even though there is very little effect on the median FWHM value, a significant increase in FWHM for the top 500 most abundant peptides is observed above 10 ng of sample load (Figure S12B). Even though data dependent acquisition settings were modified in favor of higher sample loads (TopS method), very little improvements in proteome coverage are observed when sample load is increased beyond 10 ng (Figure S12A). Further downscaling of the column cross section also has a positive effect on the elution profiles that can be achieved when working at low nano LC flow rates. With a total column volume (including connection lines) of 1.25 µL (instead of 3 µL for the micro pillar array column used in the previous publication), more efficient use of LC-MS/MS instrument time can be obtained (Figure S5C). When performing direct injection (1µL sample loop, volume is added to the total column volume) and operating the column at a flow rate of 250 nL/min, peptides start eluting as from 11.5 min after sample injection.

**References**

(1)	Stadlmann, J.; Hudecz, O.; Krššáková, G.; Ctortecka, C.; Van, G.; Beeck, J. op de; Desmet, G.; Penninger, J. M.; Jacobs, P.; Mechtler, K. Improved Sensitivity in Low-Input Proteomics Running Title : Improved Sensitivity in Low-Input Proteomics Using Micro-Pillar Array-Based Affiliations : *Analytical Biochemistry* **2019**, *91* (22), 14203–14207. https://doi.org/10.1021/acs.analchem.9b02899.

(2)	de Malsche, W.; Gardeniers, H.; Desmet, G. Experimental Study of Porous Silicon Shell Pillars under Retentive Conditions. *Analytical Chemistry* **2008**, *80* (14), 5391–5400. https://doi.org/10.1021/ac800424q.

(3)	Desmet, G.; op de Beeck, J.; van Raemdonck, G.; van Mol, K.; Claerebout, B.; van Landuyt, N.; Jacobs, P. Separation Efficiency Kinetics of Capillary Flow Micro-Pillar Array Columns for Liquid Chromatography. *Journal of Chromatography A* **2020**, *1626*, 461279. https://doi.org/10.1016/j.chroma.2020.461279.

(4)	Gritti, F.; Guiochon, G. Mass Transfer Kinetics, Band Broadening and Column Efficiency. *Journal of Chromatography A* **2012**, *1221*, 2–40. https://doi.org/10.1016/j.chroma.2011.04.058.

(5)	El-Faramawy, A.; Siu, K. W. M.; Thomson, B. A. Efficiency of Nano-Electrospray Ionization. *Journal of the American Society for Mass Spectrometry* **2005**, *16* (10), 1702–1707. https://doi.org/10.1016/j.jasms.2005.06.011.

(6)	Doblmann, J.; Dusberger, F.; Imre, R.; Hudecz, O.; Stanek, F.; Mechtler, K.; Dürnberger, G. ApQuant: Accurate Label-Free Quantification by Quality Filtering. *Journal of Proteome Research* **2019**, *18* (1), 535–541. https://doi.org/10.1021/acs.jproteome.8b00113.
